# Dynamic conformational ensembles of soluble Tau encode neuronal toxicity prior to aggregation

**DOI:** 10.64898/2026.01.26.701882

**Authors:** Florian Georgescauld, Sonia Okekenwa, Tanner Boyd, Cameron Donahue, Noe Quittot, John R Dickson, Zhanyun Fan, Scott Brady, Craig Blackstone, Bradley Hyman, Yuyu Song

**Affiliations:** Department of Chemistry and Chemical Biology, Northeastern University, 360 Huntington Avenue, MS 412TF, Boston, MA 02115; Department of Neurology, Massachusetts General Hospital and Harvard Medical School, 114 16^th^ Street, Charlestown, MA, 02129; Marine Biology Laboratory, 7 MBL Street, Woods Hole, MA 02543; Department of Anatomy and Cell Biology, University of Illinois at Chicago, 808 S. Wood St., Chicago, IL 60612

**Keywords:** Tau, Tubulin, Microtubule (MT), Hydrogen-Deuterium Exchange Mass spectrometry (HDX-MS), Protein Misfolding, Axonal Transport, Alzheimer’s Disease and Related Disorders (ADRDs)

## Abstract

Tau aggregation is a defining feature of Alzheimer’s disease and related tauopathies, yet the conformational states of Tau in neurons prior to aggregation remain poorly understood. Existing structural models are derived largely from fibrillar assemblies and provide limited insight into the dynamic, soluble Tau species that initiate pathology. Here, we combine hydrogen–deuterium exchange mass spectrometry with super-resolution imaging and neuronal models to define the conformational ensemble of soluble Tau under physiological and disease-relevant conditions. We show that soluble Tau populates distinct, dynamic conformations characterized by regional stabilization and long-range intramolecular interactions that are invisible to fibril-based structures. Disease-associated perturbations selectively remodel these conformational ensembles, exposing aggregation-prone regions and altering Tau subcellular organization in neurons. Notably, these Tau species inhibit axonal transport, which is essential for neuronal health, linking specific ensemble states to neuronal toxicity. These findings establish soluble Tau conformation as a dynamic, regulatable state that precedes aggregation and encodes disease relevance. By defining the structural logic of Tau before fibril formation, this work provides a framework for understanding early tauopathy mechanisms and for targeting Tau pathology at its earliest stages.

**SUMMARY:** Tau pathology is a hallmark of Alzheimer’s disease (AD) and related dementias (ADRDs). Although Tau is often described as intrinsically disordered, it is a dynamic protein with distinct but poorly defined conformations. Here we conduct a systematic time-resolved structure-function analysis of normal and pathologic Tau, including hyperphosphorylated, mutant Tau, and posttranslational-modification-mimetic Tau. To characterize dynamic conformational changes of Tau, we combined state-of-the-art hydrogen deuterium exchange mass spectrometry with structured illumination microscopy, demonstrating a novel Tau-MT binding mode: “dynamic oscillation”. To correlate Tau structure with neuronal function, we evaluated axonal transport as a sensitive readout of neuronal health. Many toxic Tau forms share a common signature of increased exposure of the N-terminal phosphate activating domain (PAD) *in vitro* and *in vivo*. Aberrant exposure of PAD correlates with Tau pathology and axonal transport defects. Tau phosphorylation at S262 alone is sufficient to alter Tau-microtubule interactions beyond R1-R4 motifs, globally changing Tau conformation, disrupting “dynamic oscillation” on MTs, and inhibiting axonal transport. Frontotemporal dementia-associated P301L-Tau remains associated with microtubules but also inhibits axonal transport. Our results reveal a well-defined conformation of soluble WT Tau in neurons and its highly dynamic interaction with microtubules, altered by AD/ADRD-Tau forms. Our multidisciplinary approach comprising biochemical manipulations, innovative MS tools, advanced microscopy, cellular assays, and mouse and human data pair Tau conformations with distinct neuronal functions and pathologies in health and disease.

## INTRODUCTION

A consequence of population aging has been a dramatic increase in diverse and devastating neurodegenerative diseases characterized by cognitive decline and neuronal loss^1,2^. A unifying hallmark across a major group of patients with Alzheimer’s disease (AD) and related dementias (ADRDs) is the presence of insoluble Tau aggregates within neurons, known as neurofibrillary tangles (NFT), in stark contrast to its physiologic soluble state^3–5^. However, aberrant Tau conformations, observed in Tauopathies of AD, Pick’s disease, Progressive Supranuclear Palsy (PSP), and corticobasal degeneration (CBD) present a high degree of heterogeneity in terms of cleavage^6^ and post-translational modifications (PTMs)^7,8^, as well as different fibrillar conformations^9^ and seeding properties^10,11^. This variability challenges our understanding of disease mechanisms and development of therapies. It is crucial to determine how changes in conformation and function of Tau lead to toxicity, especially before its irreversible tangle state, and how misfolded Tau leads to neurodegeneration.

Initially characterized as a microtubule-associated protein (MAP) important for microtubule (MT) assembly and dynamics, studies of Tau have revealed additional functions^12^, and the traditional view of Tau as primarily a MT stabilizer is challenged^13^. Tau is an abundant protein^14^, and MT bound vs. unbound Tau may interact with different partners and localize to various subcellular compartments^15^, where it may serve diverse roles in health and disease. For example, Tau can directly or indirectly influence other MAPs, regulating neuronal signaling pathways, cytoskeletal structures and functions. Tau can also interact with lipid membranes and may regulate synaptic vesicle and receptor trafficking, modulating synaptic function.

The capacity of Tau to form specific complexes with MTs is crucial for many of these functions, but the precise structural details and biophysical nature of these interactions remain unclear. Disease-associated Tau proteins misfold and may have altered binding to MTs, while dissociation of Tau from MTs can facilitate Tau misfolding and aggregation in solution, creating a vicious cycle. However, the mechanistic links between these dynamic processes are challenging to identify, even though they may be important for both normal function and pathology and may facilitate early diagnosis of Tauopathies. This is partly due to the difficulty in characterizing soluble Tau molecular conformations (apo or in complex) by classical structural approaches. This reflects both the lack of traditional structural motifs such as alpha helix or beta sheet, leading to the frequent description of Tau as an intrinsically disordered protein^16–21^, as well as its dynamic and transient conformations^18–22^. Recent cryo-EM studies of fibrillar Tau from patient brains^9,23^ and solid-state NMR studies of Tau in complex with MTs^16^ have elegantly defined unique Tau misfolds in disease-specific tangles and MT binding repeats (MTBRs) of Tau when bound to MTs, respectively. However, these studies have focused on specific domains of tau and are limited in their study of dynamics. Conformations of full-length Tau in either soluble pre-aggregating forms or bound to MTs remain elusive. Furthermore, questions remain about how AD/ADRD-associated mutations and PTMs of Tau^8^ induce conformational changes and affect binding to MTs, which are themselves heavily modified^24–28^. The dynamics of Tau conformation and how such changes lead to a toxic gain of function and subsequent neurodegeneration remains a puzzle. Here, we used state-of-the-art Hydrogen Deuterium Exchange (HDX) Mass Spectrometry (MS) to probe both WT and various AD/ADRD soluable Tau proteins in the presence and absence of MTs to identify dynamic structural signatures of all Tau regions. We then validated one of these conformational changes, the dynamic exposure of the N-terminal phosphatase activating domain (PAD, AA2-18), *in vivo* using mouse and human brain tissues.

To correlate conformational dynamics with function, we used axonal transport as a sensitive readout of neuronal health. Previously, we showed that most pathological Tau proteins impair axonal transport by exposing a specific region at the N-terminus, known as PAD due to its function as a phosphatase activator^29–32^. While normally sequestered, PAD’s controlled exposure activates a signaling pathway involving phosphatase 1 (PP1) and glycogen synthase kinase 3 beta (GSK3β), whose activation phosphorylates kinesin light chains and releases its cargoes for timed delivery to specific locations. Here, we hypothesized that AD/ADRD-related Tau mutations or PTMs (e.g., phosphorylation, acetylation) result in uncontrolled PAD exposure, disrupting neuronal signaling and axonal transport and triggering dying-back degeneration. To evaluate the extent to which conformational dynamics in pathological forms of Tau affect PAD exposure, we examined human full-length 2N4R Tau mutants mimicking hyperphosphorylated or acetylated Tau in AD and mutant Tau in FTD using HDX-MS and functional assays. PAD exposure increased in AD/ADRD Tau proteins, which impair axonal transport through a mechanism that was blocked by a specific antibody against the PAD motif. We further examined Tau conformations and interactions with MTs in squid axoplasm, mouse brain, and postmortem human tissue using advanced microscopic methods. By analyzing mechanisms of PAD exposure, Tau phosphorylation, and Tau-MT interactions as well as their links to Tau pathology using complementary *in vitro* and *in vivo* models for biophysical, advanced imaging, and cellular assays, we provide unique insights into the structural basis of Tau toxicity and potential therapeutic interventions.

## RESULTS

Tau is a neuronal protein with six isoforms generated through alternative splicing of the single gene *MAPT*. Structurally, it is composed of several distinct domains (see Fig. 1A for domains in the longest human isoform 2N4R): the acidic N-terminal projection domain (NPD, AA 1-165), the basic Proline-rich region (PRR, AA 166-242); the Microtubule-binding repeats (MTBRs, AA 243-367) and the C-terminal domain (CTD, AA 368-441)^12^. Each domain contains distinct subregions: PAD (AA 2-18), inserts N1 (AA 45-74) and N2 (AA 75-104) for NPD; subdomains P1 (AA 166-198) and P2 (AA 199-242) for PRR; imperfect repeats R1 (AA 243–273), R2 (AA 274–304), R3 (AA 305–335) and R4 (AA 336–367) for MTBRs; pseudo-repeat R’ (AA 368-399) for CTD. Structural studies of Tau *in vitro* showed the lack of a single stable conformation and instead existence as an ensemble of different and interchangeable conformers^33,34^. Functions have been identified for some subdomains (e.g., MTBRs), but others are defined only by exon structure in the gene or distinctive primary sequence.

**Figure 1.**
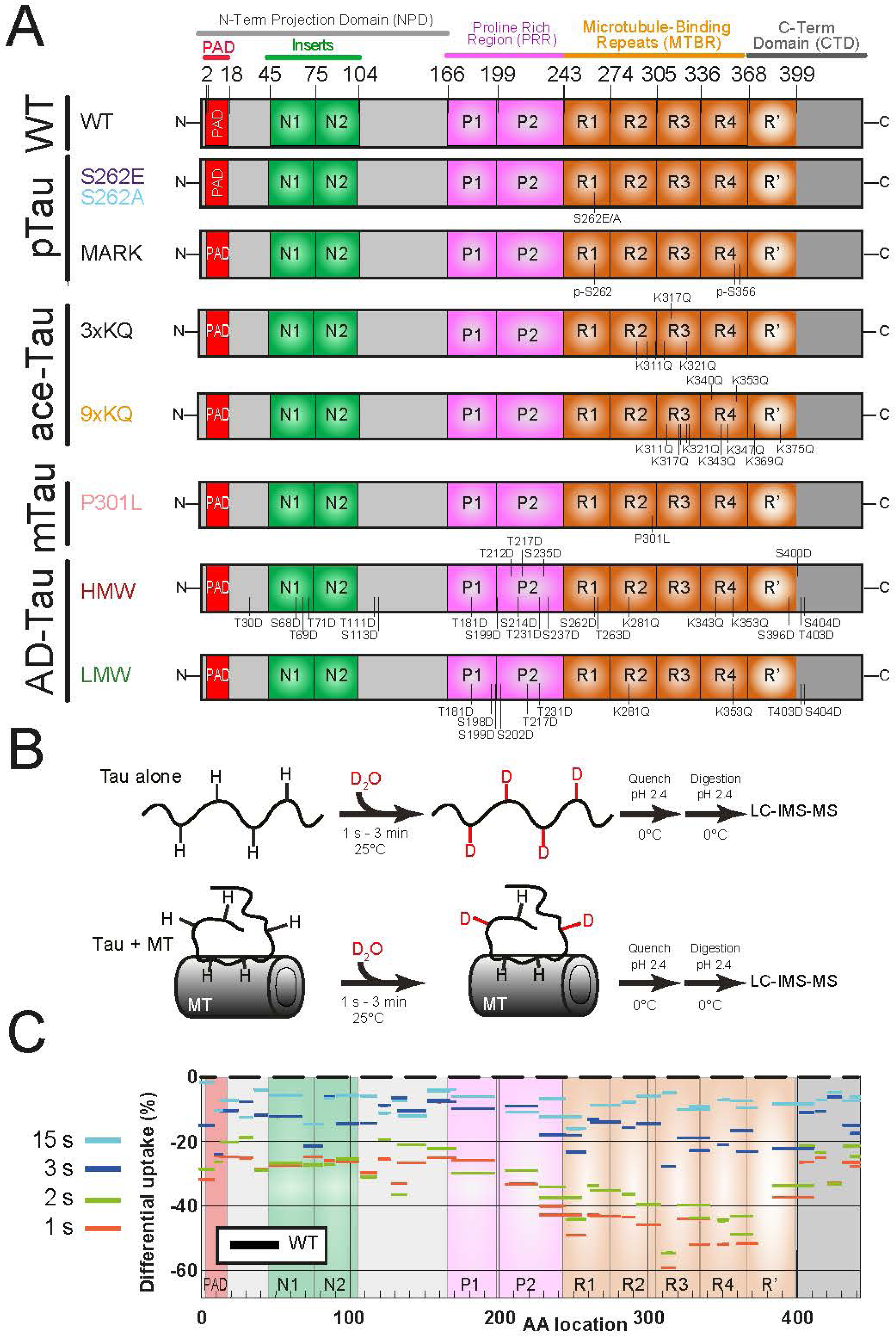
Interaction of WT-Tau with MTs as revealed by HDX-MS. (**A**) Schematic presentation of the different domains of WT 2N4R Tau (top) and various 2N4R Tau variants described in this manuscript. Each individual construct is labeled on the left side of the figure, mutations are indicated as lines (see proteins AA sequence in Figure S1A). (**B**) Schematic presentation of the HDX-MS continuous labeling experiment. Tau alone or in complex with microtubules was incubated into D_2_O buffer. Aliquots were taken at different timepoints from 1 second to 3 minutes, deuteration was stopped by acid quench on ice, and each sample was submitted to LC-IMS-MS analysis. (See Methods for details). (**C**) Relative deuteration differences for WT Tau in complex with MTs minus Tau alone, during individual deuteration timepoints (1sec-3min). Relative deuteration was expressed as percentage: 100x[Deuteration (Tau_bound_MT) – Deuteration (Tau_alone)]/(Tau full deuteration). Full deuteration was considered as the amount with Tau_alone after 3 min deuteration. Such relative deuteration indicates the protection of Tau due to MTs. As expected, MTBRs (R1, R2, R3, R4, and R’) had the least deuteration, due to direct binding to MTs. Interestingly, the protection of the rest of Tau, including N-terminal PAD motif, were also substantial, suggesting self-folding of Tau upon MT binding. Tau-MT interaction was highly dynamic, indicated by the loss of protection within seconds.

Tau is highly dynamic structurally and diverse functionally, challenging conventional protein structure-function analysis. Tau is often described as an “unstructured” protein and belongs to a group of structurally and functionally diverse proteins commonly referred to as Intrinsically Disordered Proteins (IDPs). Such proteins lack well-defined structural elements such as alpha helix or beta sheets and are typically conformationally dynamic. However, based on their biological functions, IDPs such as Tau have distinct conformations that are important for their functions. IDPs play important roles in cell signaling and structural scaffolding and are susceptible to various posttranslational modifications (PTMs). Tau is a prototypical IDP with extensive PTMs, important for cell signaling and scaffolding. Mutations and PTMs of Tau can lead to additional conformational changes, alter protein-protein interactions, and affect signaling pathways^35^. Binding to MTs dramatically restricts this dynamism, constraining Tau to a limited set of elongated conformations with unique, complementary interfaces tailored to the MT surface.

Fully characterizing the dynamic Tau-MT interaction at the three-dimensional molecular level remains exceptionally challenging due to several key factors: the inherent flexibility of Tau even when bound to MTs; the transient nature of Tau-MT interaction; and the existence of several isoforms for tubulins and Tau, both of which are subject to PTMs. This lack of detailed structural information hinders our understanding of the role of Tau in health and disease. Initial biochemical attempts on purified fragments of Tau and MTs showed that the MTBRs played a major role in binding to MTs^33^. Recent breakthroughs using cryoEM showed in detail how R1-R2 repeats interact with MTs^36^, while elegant NMR studies and single molecule fluorescence characterized stable interactions between flanking domains of the MTBRs and MTs^16,19^. Together with molecular dynamics analyses, these studies suggested that Tau interactions with MTs involved more extensive regions of Tau than MTBRs alone^37^. However, experimental data comparing pre-aggregation structures of full-length WT Tau and AD/ADRD-associated mutants or PTM modified Tau, either with or without MTs are limited. This missing gap must be filled if we want to understand how Tau folds properly to bind physiological partners (e.g., MTs) and perform normal functions in health, or misfolds to form pathological tangles and drive neuronal dysfunction in disease. Our general hypothesis is AD/ADRD Tau may alter MT binding, acquire aberrant structures, and impair neuronal function, leading to NFT formation and neurodegeneration.

### Tau Monomers are Intrinsically Disordered

To answer these questions, we cloned, expressed and purified from *E. coli* 2N4R human Tau mutant proteins mimicking those found in AD/ADRD (Fig. 1A) and the wild-type control. These modifications include a Tauopathy-related mutation: P301L, which produces frontotemporal dementia (FTD)^38,39^; single-site pseudo-phosphorylation: S262E, which mimics phosphorylated S262 that is positively correlated with clinical progression in AD^8,11,40–42^ and its nonphosphorylatable control S262A; as well as multiple-site pseudo-phosphorylation and acetylation mimicking PTMs enriched in AD patient brains^7,8^. These include 9KQ (glutamine substitution mimicking acetylated lysines), HMW (mimicking PTMs enriched in high molecular weight, seeding competent, AD-Tau) and LMW (mimicking PTMs enriched in low molecular weight, seeding incompetent, AD-Tau, also present in non-AD aged controls) (see Fig. S1A and Suppl. Table 1 for the sequence alignment and individual AA sequences). All expressed proteins were soluble. The last step of purification was size-exclusion chromatography and only monomers free of aggregates and unable to bind Thioflavin T (a fluorescent probe for amyloids) were processed and characterized by hydrogen deuterium exchange (HDX) coupled to Mass Spectrometry (MS).

Conformational dynamics for Tau monomers make determination of Tau structures challenging. HDX-MS uniquely probes protein dynamics in solution^43–46^. Solvent-accessible protein backbone amide protons readily exchange with deuterons in D_2_O within milliseconds^47^, while those buried within the protein core, involved in secondary structures or located at protein-protein interfaces, such as Tau bound to MTs, exchange much slower. Analyzing the mass shift of peptides after deuteration reveals the location and extent of hydrogen exchange, providing valuable insights into protein dynamics as well as identifying interfaces for protein complexes. We first evaluated the HDX-MS profile of Tau alone for WT and six mutants (see AA sequences in Fig. S1A) at the peptide level, according to the scheme (Fig. 1B). Each individual protein was incubated in D_2_O buffer, and aliquots were withdrawn at defined time points (1 second to 3 minutes). The deuteration reaction was quenched on ice under acidic conditions, and aliquots were treated with a protease for digestion of Tau into peptides. Deuterium incorporation into individual peptides was measured by liquid chromatography-ion mobility separation-mass spectrometry (LC-IMS-MS). High-quality peptide coverage (∼90%) was obtained for each Tau sequence (WT and mutants) (Fig. S2). Experiments were performed at least in duplicate (Suppl. Table 2).

**HDX-MS analysis of WT Tau alone** revealed complete deuteration of all identified peptides within a tight 5-10 sec window. This finding, characterized by a rapid non-sigmoidal and general plateau across all Tau regions (black curves in Figs. S1B, S3), clearly indicates the absence of significant structural barriers to exchange. Notably, for practically all peptides except for AA68-81, the maximal difference in deuteration between the plateau and the shortest deuteration time point was typically around 1 Da. AA 68-81 showed a sigmoid shape before reaching full deuteration (plateau), suggesting a rapid and transient oligomerization as previously proposed^48^. These results align with previous HDX-MS studies demonstrating ≥90% deuteration of WT Tau peptides within seconds^49,50^. Similar analysis of **point mutants S262A, S262E, P301L** showed comparable rapid deuteration kinetics, reaching complete exchange within 5-10 seconds (black curves in Figs. S1B, S4-6) and demonstrated that the mutations did not significantly alter the intrinsic disorder of Tau alone. Taken together, our results demonstrate that all regions of apo-Tau (wild-type and mutants) lack stable secondary or tertiary structures, in full agreement with the established understanding of Tau as an IDP^18,34,51^.

### Tau-MT Interactions are Dynamic

Following analysis of Tau alone, we investigated Tau **interactions with MTs using HDX-MS**. MTs were polymerized from pig brain tubulin with or without Taxol and then mixed with Tau at a stoichiometry resembling the physiological ratio in axons (5 tubulin dimers to 1 Tau monomer) till equilibrium^52,53^. Tau-MT complexes were subjected to continuous HDX-MS analysis (Fig. 1B). The different mutations of Tau and the addition of MTs resulted in distinct digestion patterns with common and variant-dependent peptides (see peptide maps in Fig S2A-G). We started the HDX analysis focusing on six peptides representative of functional regions of WT Tau (Fig. S1B). Deuteration of peptides is a sensitive probe of local environmental changes for the main chain and binary comparisons showed that deuteration profiles for Tau-MT complexes differed significantly from those observed for Tau alone. They all presented significant degrees of protection from deuteration, varying from 1.5 Da up to 5 Da at the initial 1s time point and gradually diminishing over time to reach a plateau that is similar to complete deuteration of Tau alone. This finding indicates a distinct structural rearrangement of Tau upon MT binding, with key regions within the MTBRs exhibiting the slowest deuteration, suggesting a critical role in MT interactions. Slower deuteration was also observed in the N and C termini of Tau, indicating these regions are also affected by interactions between Tau and MTs, though more transiently than the MTBRs. Similar but not identical behaviors were observed for Tau point-mutants, consistent with the sensitivity of HDX-MS in detecting dynamic interactions. To describe dynamic Tau-MT interactions in detail, we compared the deuteration patterns of each Tau variant over time and infer Tau structural changes due to MT binding from the differences in deuteration (D).

#### WT Tau with MTs

Binary comparisons for all 36 peptides of WT Tau (Fig. S3) revealed protection at various degrees across the board. To delve deeper into structural details and facilitate comparisons between peptides from each Tau variant, we converted raw deuteration values from Daltons to % relative uptake calculated as [D_(Tau + MTs)_ - D_(Tau alone)_]/D_max_ *100 (D_max_ = deuteration at the plateau). Data was plotted for each time point (Fig. 1C), with 92.8% sequence coverage (Fig. S2A). Strikingly, HDX-MS analysis painted a more nuanced picture of Tau-MT interaction/folding than traditionally understood, as protection at short times existed for all peptides, beyond MTBR, suggesting a compact Tau structure when bound to MTs. The kinetics of protection differed between peptides, as complete deuteration was reached at 5-10s for both N- and C-termini vs. 15s-1m for the MTBRs, suggesting different interaction modes consistent with Tau intramolecular interaction between N- and C-termini vs. Tau-MT intermolecular binding *via* MTBRs.

To better understand these interactions, we focused on the deuteration at time 1s when all peptides had the most protection. By plotting the 1s deuteration over the Tau AA sequence (Fig. 2G-I, black), three major regions emerged: the N-terminal region, the core-binding region, and the C-terminal region (Table 1). Detailed analyses revealed that the core binding region with the most protection (40-60% uptake difference, AA 226-372) encompasses the entire R1-R4 (AA242-368) and notably extends beyond the MTBD to include flanking portions of the proline-rich domain P2 (AA 226-241) and to some extent the R’ domain (368-372), indicating these domains constitute additional interaction sites with MTs. Based on the differences in the degree of deuteration, we divided the core binding region into five subdomains (226-243, 244-257, 258-307, 308-317, 318-372). We speculate that differences in the degree of deuteration among subdomains indicate different binding affinities and interaction lifetimes within the MT binding interface. Our HDX data showed that subdomains AA 244-257 and 308-317, which are homologous within repeats R1 and R3, stood out for very high protection. CryoEM map at high resolution only existed for complexes of MTs with the Tau repeat R1 but not R3 and showed that R1 interacts with alpha tubulin subunit^36^. The cryoEM map available for full length Tau bound to MTs at low resolution showed Tau as a continuous stretch of density along the protofilaments when covering the alpha-tubulin, but as a discontinuous one for the beta tubulin^36^. We interpret the strong protection of the HDX data for these 2 subdomains as indicative of their interaction with alpha tubulin subunits. Our % uptake data is in full agreement with this structural observation, and our findings expand the concept of Tau-MTs interaction beyond the canonical MTBD. In addition, the seemingly distant N-Terminal region (AA 8-196, 15 peptides) and C-Terminal region (AA 400-441, 5 peptides) exhibited similar, yet lower, protection (∼25% difference). Despite their sequence separation by 205 AA, they shared comparable structural behavior and interacted with neighboring segments, albeit less stably than the MTBRs. Notably, the peptide covering the first 8 residues of the PAD at the very N-terminus exhibited higher protection than the rest of the N-terminal peptides, implying a unique and distinct interaction with MTs and/or with the C-terminal of Tau as previously proposed in the paperclip model^54^. The core binding region is linked to the N- and C- terminal regions by two segments covering AA 203-225 and AA 382-399, respectively (Table 1). These segments exhibited intermediate protection and likely acted as hinges, facilitating Tau’s dynamic behavior on the MT surface.

**Figure 2.**
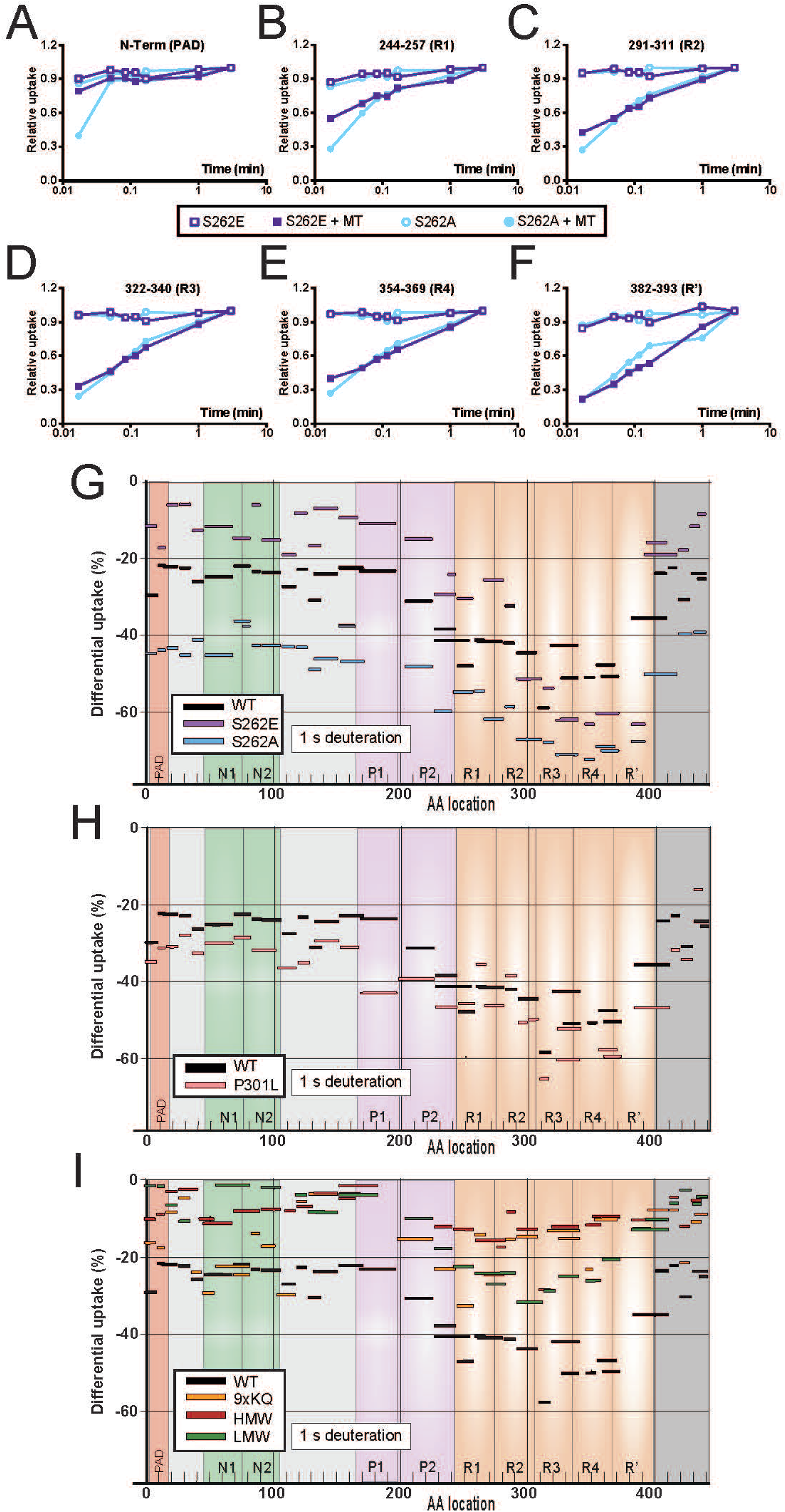
Effect of Tau mutations on its folding and interaction with MTs. (**A-F**) Deuterium incorporation with time for domain-representative Tau peptides in the absence (open symbols) and presence of MTs (full symbols) for Tau mutants S262E (purple), which resembles pathological Tau phosphorylated on serine 262, and S262A (blue), which is the non phosphorylatable control. The raw data are shown (duplicates averaged per time-point) after DynamX analysis of all common charge states followed by manual inspection and validation. (**G-I**) Skyline plots of the protection of Tau due to binding to MTs, as measured by HDX-MS at the peptide level. The position of AA of Tau variants is indicated below the graph (for individual AA mapping see Figure S3A-G) and each individual peptide of Tau is shown as a stick. The protection measured at 1 second is indicated on the Y axis as % deuteration values. For each individual peptide, the protection was calculated by measuring: 100x[Deuteration (Tau_bound_MT) – Deuteration (Tau_alone)]/(Tau full deuteration). Tau different variants are: (**G**) WT (black), S262A (blue) and S262E (magenta); (**H**) WT (black) and P301L (pink); (**I**) WT (black), 9xKQ (orange), LMW (green) and HMW (dark red).

Tau was discovered because it copurifies with MTs, yet the nature of MT binding is not fully understood. Our HDX data shows a differentiation of subdomains within the core binding region, along with the presence of hinge segments flanking it, which is indicative of a “*continuous oscillation*” of Tau on the MT surface with rapid binding-release alternating between different Tau segments. We propose that individual subdomains within the core region detach from the MTs in a transient manner while the neighboring fragments remain bound, maintaining Tau’s overall proximity to the MTs. This dynamic interaction significantly prolongs the overall association of Tau with MTs while explaining the complete deuteration of its peptides within a short time frame.

#### WT Tau with MTs stabilized by Taxol

MT depolymerization, although unlikely at equilibrium above the critical tubulin concentration and warm temperature during these experiments, could potentially contribute to observed loss of protection of Tau against deuteration over time. To rule out this possibility, we added paclitaxel (Taxol) to preassembled MTs. Taxol does not directly interact with Tau and is a widely used anticancer drug, which stabilizes MTs by hindering tubulin conformational changes and structurally inducing a GTP-like state^55^. Compared to previously analyzed Tau-MT complexes, Taxol induced a significant reduction (∼20-25%) in the protection against deuterium exchange across all Tau peptides (Figs. S7 & S8). This data suggests that stabilization of preassembled MTs and potential alteration of the MT lattice by Taxol may decrease Tau-MT binding and/or self-folding. This supports our assumption that MT depolymerization doesn’t contribute to increased deuteration of Tau observed. Taxol binding to MTs leads to a more transient and weakened Tau-MT binding mode, perhaps due to Taxol-induced MT conformational changes that affect the interaction surface on tubulin for Tau and/or direct competition for binding sites. For the previously described 3 major regions of Tau, the core binding region (AA 226-372) retained moderate protection, while the N-terminal (AA 1-196, including the PAD) and C-terminal (AA 400-441) regions, which showed ∼25% protection without Taxol, became rapidly deuterated. This suggests a conformational change that affects Tau folding/binding and may lead to a gain of toxicity in structurally open Tau. This finding may also explain, at least partially, why MT stabilizing drugs have been unsuccessful in clinical trials for AD and spinal cord injuries. Finally, we found that the two hinge segments exhibited differential responses to Taxol. The N-terminal hinge (AA 203-225) showed minimal protection, while the C-terminal hinge (AA 382-399), corresponding to the R’ domain, displayed a similar degree of protection to that of R1 and R2 (AA 382-410 and AA 400-410). This suggests a compensatory strengthening of its MT binding in response to the weakening of Tau-MT affinity in other domains and indicates a more prominent role for the R’ domain in stabilizing Tau binding to Taxol-stabilized MTs, consistent with NMR data using similar conditions^16^.

#### S262E pseudophosphorylated Tau with MTs

Residue S262 is universally conserved among Tau isoforms. S262 phosphorylation was proposed to attenuate MT binding and to correlate with AD progression in patients^11^. We studied mutant S262E Tau which mimics such phosphorylation. The S262E Tau HDX profile differed significantly from WT Tau (purple skyline plots in Fig.2, kinetics of representative peptides in Figs.S1B, S4 and Table.1). The core binding region (AA 291-393, 7 peptides) encompasses the second half of repeat R2, full repeats R3 and R4, and almost the entirety of the R’ domain, and presented a remarkably high uptake difference of 52-62%, suggesting strong binding to MTs similar to that of WT. AA 322-393 (5 peptides covering the second half of R3, full R4, and most of the R’ domain) displayed the highest protection, highlighting its role as the primary contributor to MTs binding. In contrast, the N- and C-terminal regions (AA 1-225, 16 peptides and AA 392-441, 5 peptides, respectively) demonstrated weak protection (∼12% uptake difference), which was significantly lower than in WT, suggesting reduced interactions with MTs and/or self-folding. However, while these values were low, they were not zero, indicating measurable, albeit fleeting interactions. Significantly, protection of the PAD within the N-terminal region was minimal, aligning with the average values of neighboring peptides, suggesting an open and accessible nature in contrast to the putative paperclip fold in WT Tau. The core binding region connected to the N-terminal region through a “linker” formed by a five-peptide segment (AA 226-290) exhibiting intermediate protection (25-32% uptake difference). No comparable “linker” was detected between the highly protected region (AA 291-393) and the C-terminal one (AA 392-441).

The S262E mutation resides in the heart of repeat R1 and significantly reduced protection of R1 against deuteration. Based on the structure of the R1-MT complex (PDB 6CVJ), S262 directly faces tubulin residue E434. Mutation of S262 into E results in electrostatic repulsion and destabilization of overall interaction as we determined by calculating the Electrostatic Potential Surfaces for the mutant and comparing it to the WT (Fig.S1C). Surprisingly, this destabilization propagates across Tau’s entire sequence, except for the stretch between distal half of R2 and R’, as revealed by comparing WT and S262E (Figs 2G & S1B). First, the segment encompassing the mutation (AA 226-290) belongs to the core binding region in the WT but transforms into a “linker” with intermediate protection in the mutant. This implies the mutation disrupts Tau’s binding mode within this crucial region. Second, the N-terminal region (AA 1-226) exhibits significantly weaker protection in the mutant compared to WT, indicating more fleeting interactions with potential partners. This altered behavior extends beyond the immediate vicinity of the mutation. Third, the most protected core binding region becomes significantly shorter in the mutant (AA 291-393) compared to WT (AA 226-372) and features a partially different sequence composition, with the R’ domain now integrated within the core binding region instead of serving as a linker in WT. Interestingly, despite the shorter core binding region (102 AA in mutant versus 146 AA in WT), this region of S262E mutant displayed higher uptake difference values, suggesting stronger affinity to MTs locally, potentially as a compensatory mechanism. This enhanced binding could be attributed to the R’ strong MT interaction, which in turn eliminates the moderate protection of the linker region observed in WT and induces weaker protection in the C-terminal region. In conclusion, although the S262E mutation is located in the center of the Tau sequence, it exerts profound and unexpected effects across the entire protein sequence while bound to MTs. This highlights the intricate interplay between various Tau domains and the far-reaching consequences of seemingly localized modifications and points to a high degree of structural cooperativity within Tau. We predict that changes in one domain (due to the mutation or PTM) can ripple through and influence the conformation and dynamic behavior of other more distant domains, like the N- and C-termini.

To determine whether the Tau S262E mutant could present a “continuous oscillation” along MTs like the WT, we focused on the core binding region. Interestingly, its first two peptides (AA 291-321) showed similar protection, while the remaining five (AA 322-393) exhibited remarkably consistent and stronger protection. This pattern suggests that these five peptides act as a unique structural unit, simultaneously bound or unbound to the MTs. Unlike the WT Tau, which presents 5 subdomains within its core binding region surrounded by two linkers of intermediate protection, the presence of only 2 of them with a single linker for the S262E mutant implies that it cannot present a “continuous oscillation” on the MT but instead detaches as a whole molecule.

Altogether, these findings emphasize the important effects of the S262E mutation on Tau-MT interactions and Tau self-folding, revealing a shift from dynamic interactions of compactly folded WT Tau in proximity to MTs to potentially complete detachment of loosely open mutant Tau and highlighting the intricate interconnectedness of Tau structural domains.

#### S262A non-phosphorylatable Tau with MTs

S262A Tau exhibited the most extensive protection across all its peptide segments compared to WT and S262E (blue plots in Fig.2, and Figs.S1B, S5). This heightened protection existed in three distinct regions (Table 1). The most prominent feature was the expanded core binding region, encompassing residues 226-393 (13 peptides) and serving as the anchor point for MT binding. This region could be further dissected into two sub-fragments: the stronger binder (AA 291-393) comprising half of R2, full repeats R3 and R4, and the R’ domain, and the slightly less pronounced binder (AA 226-290) including half of P2, full R1, and half of R2. Flanking this central stronghold were two additional protected zones, the N-and C-terminal regions (AA 8 to 225 and AA 419-441, respectively). Although exhibiting weaker protection compared to the major binding area, both regions displayed a surprisingly high percentage uptake difference of around 40%, mirroring the pattern observed in the WT Tau. Like the WT, the peptide covering PAD motif at the very N-terminus exhibited high protection, consistent with the paperclip conformation where PAD is sequestered by C-terminus. Additional linker segments bridging the N- and C-terminal regions with the core binding region existed and displayed an intermediate level of protection, further highlighting the intricate structural organization of S262A in its MT-bound state, most similar to that of the WT. Altogether, S262A Tau paints a picture of Tau-MT interaction characterized by an expanded and fortified core binding region with significant protection in terminal regions. This protection pattern is similar to the WT, though stronger, perhaps due to the smaller side chain in alanine that may fit more tightly into the MT pocket.

#### P301L FTD-Tau with MTs

The deuteration profile of P301L Tau bound to MTs revealed a distinct pattern compared to WT Tau (coral color plot in Fig.2H, and Figs.S1B, S6). The most striking feature was the expansion of the core binding region, which stretched across 244 residues (AA 167-410, 15 peptides). This region was further dissected into three large subdomains. The central fragment (AA 291-372), constituting the heart of the core binding region, displayed remarkable protection with percentage uptake difference of 50-65%. It encompassed distal half of R2, full repeats R3 and R4, and served as the primary anchor point for MT interaction. Interestingly, its behavior and protection values mirrored those observed in the S262E mutant, suggesting a similar binding mode. Based on the PDB structure, we further observed that the distal half R2 within this fragment engaged in extensive interactions with a full beta-tubulin monomer, potentially explaining its enhanced binding strength compared to the other half R2 segment. N-Terminal Fragment (AA 167-290), despite being slightly more accessible to solvent than the central fragment, still exhibited significant protection with a 35-48% uptake difference, comparable to the MTBRs of WT Tau. Notably, it contains the Proline-Rich Domains (PRDs) P1 and P2, R1, and half of R2, indicating a significant contribution of these domains to MT binding in P301L, more so than in the WT. The C-Terminal Fragment (AA 382-410), encompassing half of R’ and the beginning of the C-terminal domain, also displayed strong protection with binding to MTs, more than that of the WT.

Altogether, these findings demonstrate that P301L utilizes a significantly larger segment of the Tau sequence for MT interaction compared to WT Tau, with the strongest binding occurring through a continuous fragment containing half-R2, R3, and R4. Notably, all peptides within this fragment exhibited higher protection in P301L compared to their WT counterparts. Additionally, the N- and C-terminal regions (AA 8-166 and AA 411-436, respectively), together with a protected PAD motif, displayed slightly higher uptake difference values compared to WT Tau, suggesting potential interactions with yet-unidentified binding partners, which could occur intramolecularly or intermolecularly and will be addressed by immunofluorescence staining later. Finally, the varying binding amplitudes observed within subdomains of the core binding region indicate that P301L undergoes dynamic movements on MTs without completely detaching similar to WT.

#### Other Tau variants mimicking AD Tau

Finally, we examined acetylation mimic Tau (9KQ), as well as acetylation + phosphorylation mimic HMW Tau (HMW) in AD and LMW Tau (LMW) in both AD and aged non-AD controls. HDX-MS profiles of these variants were obtained in a similar manner (Figs. 2I, S9-11) with ∼90% peptide coverage (Fig.S2E-G).

#### 9KQ pseudoacetylated Tau with MTs

9KQ mutant displayed unique behavior compared to other mutants and WT Tau. Notably, even in the absence of MTs, some of its peptides exhibited partial protection, exceeding 1 Da in deuteration for regions including AA 1-14, 54-81, 90-101, 127-150, and 428-436 (Fig.S9). This suggests the formation of small compact subdomains within 9KQ Tau itself, with highly dynamic nature as evidenced by their complete deuteration within 10s. Size Exclusion Chromatography analysis confirmed that these subdomains did not lead to detectable aggregation throughout the duration of these experiments, as 9KQ’s elution volume remained similar to WT Tau. In the presence of MTs, 9KQ mutant displayed a distinct pattern of alternating slightly protected and unprotected peptides (orange plot in Fig.2I), unlike the usual binding regions observed in WT Tau, suggesting a complete shift in its MT binding capacity. Instead of stable interactions, weaker and transient contact points with MTs were observed, indicating a high propensity for MT detachment, opportunity for self-aggregation, and potential functional consequences on the neuronal cytoskeleton.

#### HMW pseudoacetylated and pseudophosphorylated Tau in AD

HMW mutant also exhibited a distinct set of interactions as compared to WT Tau and point mutants (red plot in Fig.2I). Notably, in the presence of MTs, none of the HMW peptides displayed the characteristic sigmoidal shape in HDX-MS analysis, which is indicative of binding to a partner molecule (Fig.S10). This observation was further corroborated by the modest protection levels observed for most peptides at deuteration kinetic 1 second, ranging from 0-15% (Fig.2I). In HWM Tau, only peptide AA 308-317, which is part of R3 with high protection (∼60% or more) in WT, S262A, and P301L Tau proteins, showed slightly higher protection (25%) than the other domains. These findings suggest that HMW Tau’s interaction with MTs is significantly weaker than the canonical Tau-MT binding, while a partial and transient interaction with MTs cannot be entirely ruled out.

#### LMW pseudoacetylated and pseudophosphorylated Tau in AD and aged controls

The HDX profile of LMW mutant (green plot in Fig.2I) in the presence of MTs resembled, to some degree, that of WT Tau with Taxol-stabilized MTs, indicating a reduced yet measurable MT binding. Notably, both N- and C-terminal fragments displayed reduced protection (Fig.S11) in the presence of MTs, indicating the tendency to lose the compact conformation observed in WT Tau. However, a transient interaction with MTs was observed for the continuous central region (AA 226-372) with an uptake difference ranging 18-32%. This region mirrors the core binding region of WT Tau. Interestingly, the protection within this region followed a V-shaped pattern, increasing from the extremities towards the center, formed by the second half of R2 (AA 291-311). Our HDX data shows that this segment acts as a major binding anchor to beta-tubulin monomer as suggested by cryoEM data (PDB 6CVK and 7PQC).

In contrast to the robust and extended MT binding observed for WT, S262A, and P301L, the introduction of multiple changes to the Tau sequence to mimic Tau PMTs in AD brains (9KQ, HMW) and in both AD and aged non-AD brains (LMW) show significantly reduced MT interaction and loss of the paperclip fold, though to various degrees. HDX-MS analysis reveals minimal to no protection for these mutants in the presence of MTs, suggesting at best fleeting and transient interactions. Further investigation reveals significantly greater exposure of their PAD motif compared to WT Tau, ranging from almost complete accessibility in HMW and LMW Tau to partial protection in 9KQ Tau. This heightened accessibility of PAD motif, which is known to interact with PP1 when aberrantly exposed and inhibit axonal transport to cause neuronal toxicity, could potentially contribute to AD pathology as will be examined in detail below. These findings highlight the diverse ways Tau mutations and PTMs impact Tau-MT interactions and Tau self-folding, warranting further investigation into their specific roles in Tau-related pathologies at various disease stages. To begin addressing the functional implications of these structural changes, we next studied the effects of these variants on axonal transport, the role of PAD and the PP1 signaling pathway, and Tau-MT interactions using a unique model: isolated squid axoplasm. We then tracked PAD exposure, Tau phosphorylation on S262, and Tau-MT interaction in THY22 mouse models^56,57^ with progressing Tau pathology (3-24mo) and postmortem human tissues with various levels of Tau PTMs.

### Inhibition of Axonal Transport by Tau Variants Mimicking AD/ADRD Tau Species

To correlate Tau structural changes with neuronal function and to study the unique gain-of function effects of Tau variants, we used isolated axoplasm from squid and measured vesicular transport rates^58^, a sensitive assay to evaluate neuronal health, because axonal transport of vesicles is essential for axonal maintenance and synaptic function. The isolated axoplasm model provides intact MT cytoskeleton and functional axonal transport machinery, both of which are highly conserved in the mammalian system, ensuring the potential of translation. When perfused with buffer X/2, a physiological buffer for axoplasm, cytoskeletal structures are dispersed to allow better visualization and quantification of MT structure and its interaction with other cytoskeletal components (e.g., Tau), as well as vesicular trafficking along MT tracks^58–60^. Our previous studies suggested that multiple pathological Tau proteins inhibited axonal transport *via* the aberrant exposure of PAD motif and downstream activation of the PP1-GSK3β pathway which inhibits kinesin motility without affecting dynein^29,32^. Here, we tested all the variants analyzed by HDX-MS and found that S262E Tau significantly inhibited anterograde transport without affecting retrograde (Fig.3A) and such an inhibition was mediated by PAD motif as sequestering PAD motif using TNT1 antibody, which selectively binds AA 7-12 of Tau^29,32^, prevented the transport defect induced by S262E Tau (Fig.3B). Since S262 site is phosphorylated by MARK kinases, which are elevated in AD, we performed *in vitro* phosphorylation of Tau by MARK2 to generate MARK-Tau, which similarly inhibited anterograde transport (Fig.3C) and was blocked by TNT1 antibody (Fig.3E). In contrast, nonphosphorylatable S262A Tau had no effect on axonal transport (Fig.3D), like the WT and buffer alone control (Fig.3E). In addition, 800nM Okadaic Acid, which inhibits PP1, fully rescued axonal transport defects caused by S262E and MARK-Tau (Fig.S12). Together with the HDX-MS data showing the exposure of PAD in S262E Tau, these data suggest that S262E Tau inhibits anterograde axonal transport *via* PAD exposure, which is known to bind and activate PPI, leading to phosphorylation of kinesin by GSK3 and reduced motility of vesicles^29,32^.

**Figure 3.**
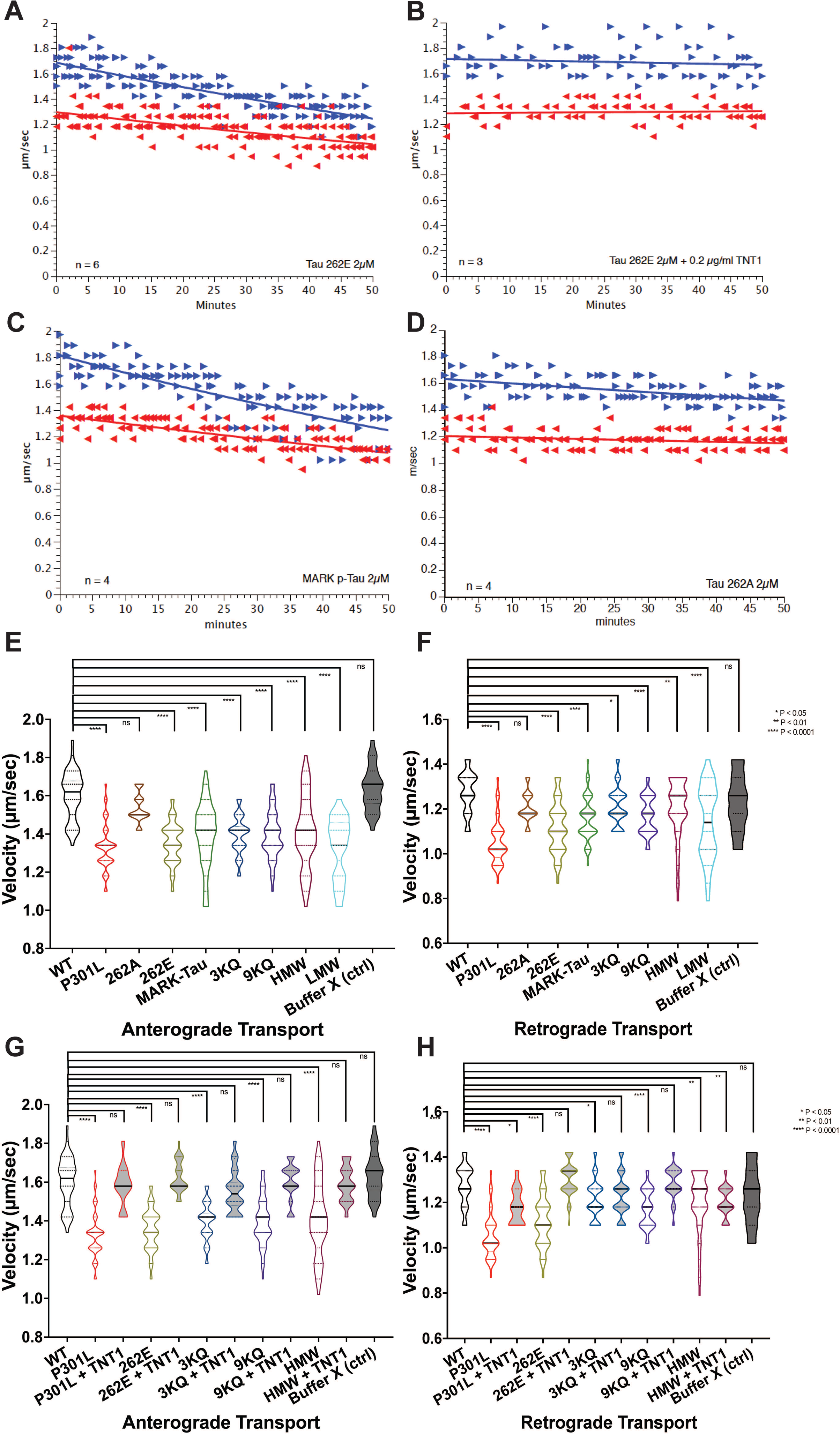
Effect of Tau mutations on axonal transport. (**A**) Phosphomimic S262E-Tau inhibited axonal transport, predominantly in the anterograde direction. (**B**) TNT1, an antibody targeting N-terminal PAD motif, prevented the S262E-induced inhibition of axonal transport. (**C**) In vitro phosphorylated Tau by MARK also inhibited axonal transport similar to S262E-Tau. (D) Nonphosphorylatable S262A, as a control, didn’t affect axonal transport. (**E-F**) Quantification and statistics of the effects of all Tau mutants on anterograde and retrograde axonal transport, respectively. (**G-H**) Quantification and statistics of the rescuing effects of TNT1 against selective Tau mutants on anterograde and retrograde axonal transport, respectively. Note that though TNT1 fully rescued anterograde transport for all Tau variants tested, it failed to rescue the retrograde transport defect caused by P301L-Tau and HMW-Tau, suggesting additional pathways independent of PAD motif.

Interestingly, FTD-P301L Tau inhibited both anterograde and retrograde transport (Fig.3E-F). While sequestering PAD using TNT1 antibody prevented P301L-induced anterograde transport inhibition as it did for S262E, retrograde transport rate was not fully recovered. Our HDX-MS data suggests full protection of PAD in P301L Tau in the presence of MTs and expanded and enhanced MT binding region, which cannot explain the transport data here. We speculate that in a saturating condition as used in HDX-MS, P301L-Tau may form intermolecular interactions that may lead to the formation of Tau envelope demonstrated by others^61^. However, in the axoplasm where added P301L-Tau:MT is 1:12.5, such intermolecular interactions may not form, so P301L Tau may remain as an open structure exposing PAD and potentially exposing other sites that could interact with different partners. Alternatively, endogenous WT Tau bound to MTs in the isolated axoplasm system may limit the MT binding sites for P301L Tau, causing P301L to detach from MTs and induce its open structure. To examine these possibilities, we performed immunohistochemistry (IHC) using TNT1 antibody that specifically recognizes PAD and confirmed that P301L Tau was bound to MTs and that PAD was exposed (Fig.S13). Given the unique HDX profile of P301L Tau when bound to MTs and the additional inhibition of retrograde transport that was not prevented by blocking PAD motif, we speculate that additional misfolded motifs and/or binding partners may contribute to the transport defects caused by P301L.

Finally, 9KQ, 3KQ (with pseudoacetylation on R3 alone), HMW, and LMW Tau all inhibited axonal transport in both anterograde and retrograde directions (Fig.3E-F). Sequestering PAD by TNT1 rescued transport defects in both directions for 9KQ and 3KQ but only rescued anterograde transport for HMW (Fig.3G-H), suggesting additional mechanisms inhibiting dynein by HMW, which is beyond the scope of current study.

### PAD Exposure and Altered MT Interactions in AD/ADRD Tau Mimics

To examine the effects of various Tau modifications on Tau structures and Tau-MT interactions in a more physiological setting, we examined squid axoplasm, where arrays of MTs and their dynamic interactions with other MAPs are naturally preserved, using an enhanced resolution imaging technique – structured illumination microscopy (SIM). In the isolated axoplasm, WT Tau was well incorporated into the endogenous axonal MTs as shown by both Atto488 labeled (Fig.4A-F) and unlabeled Tau (Fig.S13S-X), suggesting the normal capacity of recombinant Tau in binding endogenous MT protofilaments and sufficient binding sites for Tau on these MTs. PAD exposure in WT Tau was limited as detected by TNT1 immunofluorescence. Consistent with the HDX data, S262E Tau showed reduced MT co-localization, aggregation in areas where MTs were sparse, and increased PAD exposure (Fig.4G-J, S-T), while S262A behaved like WT Tau except for increased MT association (Fig.4K-N, S-T). Consistent with the prior HDX data, P301L Tau was well associated with MTs (Figs.4T, S13A-F). However, PAD was exposed in P301L Tau when perfused into the axoplasm, consistent with the transport data where sequestering PAD prevented P301L-Tau-induced anterograde transport defect.

**Figure 4.**
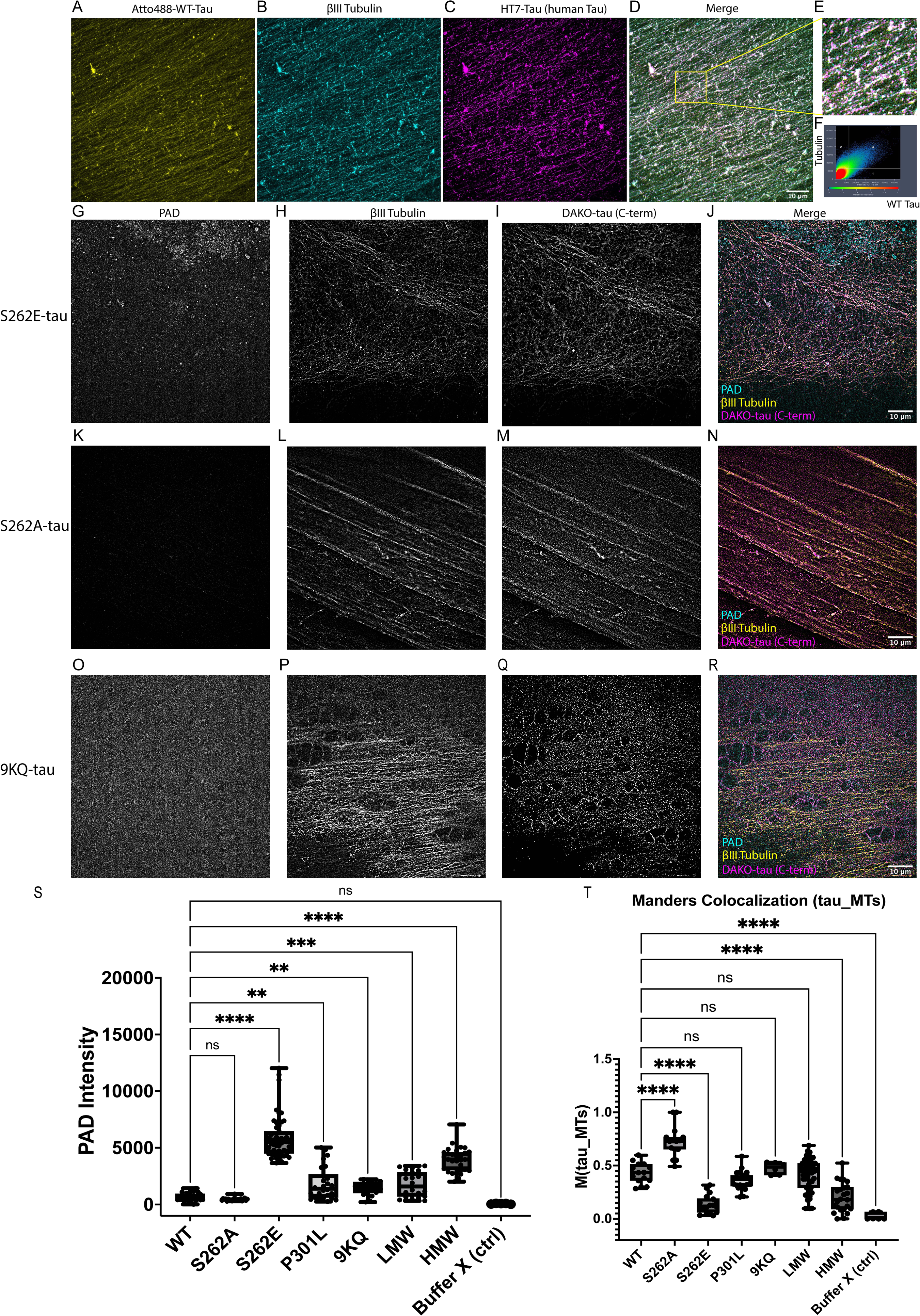
Effect of Tau mutations on axonal MT binding. (**A-F**) WT human 2N4R Tau conjugated to atto-488 perfused into the isolated axoplasm was fully incorporated into endogenous MTs and confirmed by human Tau antibody (HT7). (**G-J**) Phospho-mimic Tau with S262E point mutation was partially incorporated into MTs and showed high PAD immunoreactivity. (**K-N**) Nonphosphorylatable Tau with S262A mutation was fully incorporated into MTs, as WT Tau, and showed minimal PAD immunoreactivity. (**O-R**) Pseudoacetylated Tau with multiple K-Q mutations was also incorporated into MTs with increased PAD immunoreactivity. Interestingly, 9KQ-Tau seemed to disperse neighboring MTs and created empty spaces not seen in other experiments. (**S-T**) Quantification and statistics of PAD immunoreactivity and Tau-MT association.

Interestingly, 9KQ Tau was associated with MTs at a level similar to WT, with a diffuse pattern of moderate PAD exposure but a striking change in MT organization, which appeared as increased spacing and gaps between neighboring MTs (Fig.4O-R, S-T). Considering the unique HDX profile of 9KQ Tau, which exhibited transient interactions and compact small subdomains within the 9KQ Tau protein itself, we speculate that local self-folding of these subdomains may drive associated MTs into the circular structures seen here. As acetylation is an important PTM associated with hyperstable MTs, it will be interesting to examine in future studies whether this PTM on MTs may compensate for the MT structural perturbation caused by acetylated Tau. Finally, HMW Tau, which mimics higher order Tau oligomers enriched in AD patient, showed significantly decreased MT association and increased PAD exposure in axoplasm (Figs.4S-T, S13G-L). In contrast, LMW Tau, which models low molecular weight Tau enriched in AD but are also present in control age-matched patients, showed increased PAD exposure as compared to WT but lower than S262E or HMW Tau species (Figs.4S-T, S13M-R), and a similar level of MT association to that of WT Tau. For both HMW and LMW mutants, the SIM data was largely consistent with HDX and transport data.

In the light of the HDX data on Tau conformations and Tau-MT interactions *in vitro* and the transport data on the effects of Tau variants on neuronal function *ex vivo*, SIM imaging of Tau-MT association suggests that WT and S262A non-phosphorylatable Tau maintain a folded structure with PAD motif sequestered and a tight association with MTs, while phosphorylation on residue S262 triggered an open conformation of Tau with exposure of the PAD and release from MTs, resulting in inhibition of axonal transport and local aggregation. As more PTMs (phosphorylations and acetylations) are added to Tau, LMW Tau in aged brains tends to lose its compact fold to expose PAD and inhibit axonal transport, though still being associated with MTs. In contrast, AD-associated HMW Tau fully loses its normal folding to completely expose PAD and is released from MTs to potentially aggregate into fibrils. Acetylations alone in 9KQ don’t seem to completely dissociate Tau from MTs but alter local spacing of MTs.

P301L-Tau also seems to be well incorporated into MTs, consistent with the HDX results. Interestingly, PAD exposure is increased in both 9KQ and P301L, despite their tight MT binding, suggesting an intermediate form of open Tau that stays on MTs and impairs axonal transport by recruiting PP1-GSK3β to phosphorylate and inhibit kinesin motility. Our data demonstrates changes in Tau conformation and Tau-MT interaction caused by individual Tau variants, indicating a progressive conformational change of Tau and its dissociation from MTs during disease progression. To determine whether this is consistent with AD/ADRD pathogenesis, we turn to an FTD-Tau mouse model and human AD patient samples to evaluate PAD exposure and Tau phosphorylation, especially at S262 site, as well as MT association in the context of Tau pathology and disease progression.

### PAD Exposure, Phosphorylation of S262, and Tau-MT association in THY-Tau22 Mouse Brain

THY-Tau22 mice bear two human FTD-associated mutations: MAPT G272V and MAPT P301S and have been widely used as a model for studying Tau pathology, such as hyperphosphorylation, aggregation, neurofibrillary tangle (NFT) formation, in AD/ADRDs. These mice show 4∼5-fold increase in human Tau over endogenous Tau in the cortex at 3mo^62^, demonstrate synaptic changes as well as learning and memory deficits without motor problems at 6-9mo, and display neuronal loss at 12mo^56,62,63^. Here, we examined the two signatures of misfolded Tau in our study: PAD exposure and phosphorylation at S262 (p262), in various brain regions: Entorhinal Cortex (EC), CA1, CA3, and Dentate Gyrus (DG) of these mice at 3mo, 6mo, and 24mo. We found that PAD immunoreactivity, when normalized to total Tau level, was significantly increased (∼2 fold increase) in EC, CA1, CA, and DG of the THY-Tau22 mouse, compared with their WT littermates (Fig.S14) as early as 3mo. Interestingly, PAD exposure seemed to plateau as its immunofluorescence intensities didn’t increase in 6 and 24mo animals, suggesting that mutant Tau species in these transgenic mice intrinsically lost their paperclip self-fold long before any severe Tau pathologies, synaptic dysfunction, and behavioral impairment. In the meantime, p262 Tau, when normalized to total Tau level, was also significantly higher in all EC and hippocampal regions of the THY-Tau22 mice than the WT and continued to increase in the mutants as they aged from 3mo to 6mo, though its level decreased to the level of 3mo at the end stage (24mo), perhaps due to neuronal cytoskeletal loss, Tau conformational changes and aggregation, or/and altered signaling pathways and proteostasis. Given the HDX data and axoplasm IHC data showing decreased MT binding of pseudophosphorylated S262E Tau, we speculated that increased p262 would also release Tau from MTs. Therefore, we examined Tau-MT association using SIM imaging and SIM^2^ analysis as the next step.

Because PTMs and morphologies of both Tau and MTs differ in the soma vs. axon and Tau mutants including P301L and S262E inhibit axonal transport, a direct early axonal defect that causes dying back neuropathy in neurodegeneration, we quantified Tau-MT association in the cell body of EC vs. axons in the corpus callosum (CC) at 3, 6, 9, 12, 15, 18, 24mo (Fig.S15). Tau was highly associated with MTs in the cell body except for a few places where Tau accumulation was accompanied by faint tubulin staining throughout all age groups (Fig.S16), while Tau association with MTs in the axonal tracts was significantly reduced at 9mo and remained afterwards. Although the difference in Tau-MT association between 6mo and 3mo didn’t reach statistical significance, there was a trend toward decreased binding at 6mo, consistent with the increase in p262 Tau level observed earlier.

Altogether, these data suggest that Tau in this FTD mouse model has an intrinsic open structure exposing PAD motif at 3mo or earlier, with significant loss of Tau bound to MTs in axonal tracts with the earliest trend at 6mo, corresponding to the time frame of p262 increase in these mice. Given the pathogenicity of PAD in inhibiting axonal transport, these misfolded Tau species are likely to cause the synaptic dysfunction and memory loss observed later. It is also worth noting that despite a general progressive pattern, not all neurons are similarly affected in the mutant mice at each stage (3-24mo), some neurons seem to show higher Tau accumulation accompanied by increased PAD immunoreactivity even at an early stage while some neurons show decent Tau-MT association and relatively normal morphology at the final stage. Similarly, some axons seem to lose Tau association early on while others are preserved at the end stage. This heterogeneity within the same brain region suggests selective vulnerability of some neurons and axonal tracts with significant Tau accumulation and loss of MT binding. Equally interesting is the population of neurons and axons that seem to be relatively spared. Similar heterogeneity has also been observed in human patients where Tau PTMs and seeding competence vary from patient to patient^7,8,10,11^. Therefore, as a final step, we turned to representative postmortem AD patient tissues with well described Tau pathologies to study PAD exposure, p262 level, and Tau-MT association in the disease context.

### PAD Immunoreactivity, Tau Phosphorylation on S262, and MT Interaction in AD Patient Hippocampus

Consistent with previous reports^29,32^, PAD immunoreactivity was much higher in the hippocampus of the AD patients, compared with age/sex/PMI-matched controls, and Tau tangles were also positive for PAD (Fig.5 A-F, M). Consistent with the mouse data, p262 Tau level was also significantly higher in AD hippocampus (Fig.7 G-L, N), following a similar pattern as PAD staining. Interestingly, dot blots of HMW Tau fractions from 7 patients previously profiled for PTMs using proteomics and evaluated for seeding competence using biochemistry^7,8^ showed a trend of correlation where the highest PAD and p262 immunoreactivities corresponded to the AD cases with the most PTMs (including phosphorylation, ubiquitination, and acetylation) and highest seeding activities (Fig.7 O-P). When studied in more detail, we found that a higher heterogeneity and a similar progressive pattern existed in AD patients, compared with the mouse data. For example, in Braak III-IV cases, despite the high PAD exposure, PAD immunoreactivity in most neuronal cell bodies was still spatially associated with MT distribution, though this association was significantly lower in the axonal tracts (Fig.S17). However, at Braak VI where PAD immunoreactivity remained high, its co-localization with MT was largely absent in both neurons and axonal tracts and this was confirmed using multiple enhanced resolution imaging techniques, including STED imaging (Fig.S18).

**Figure 5.**
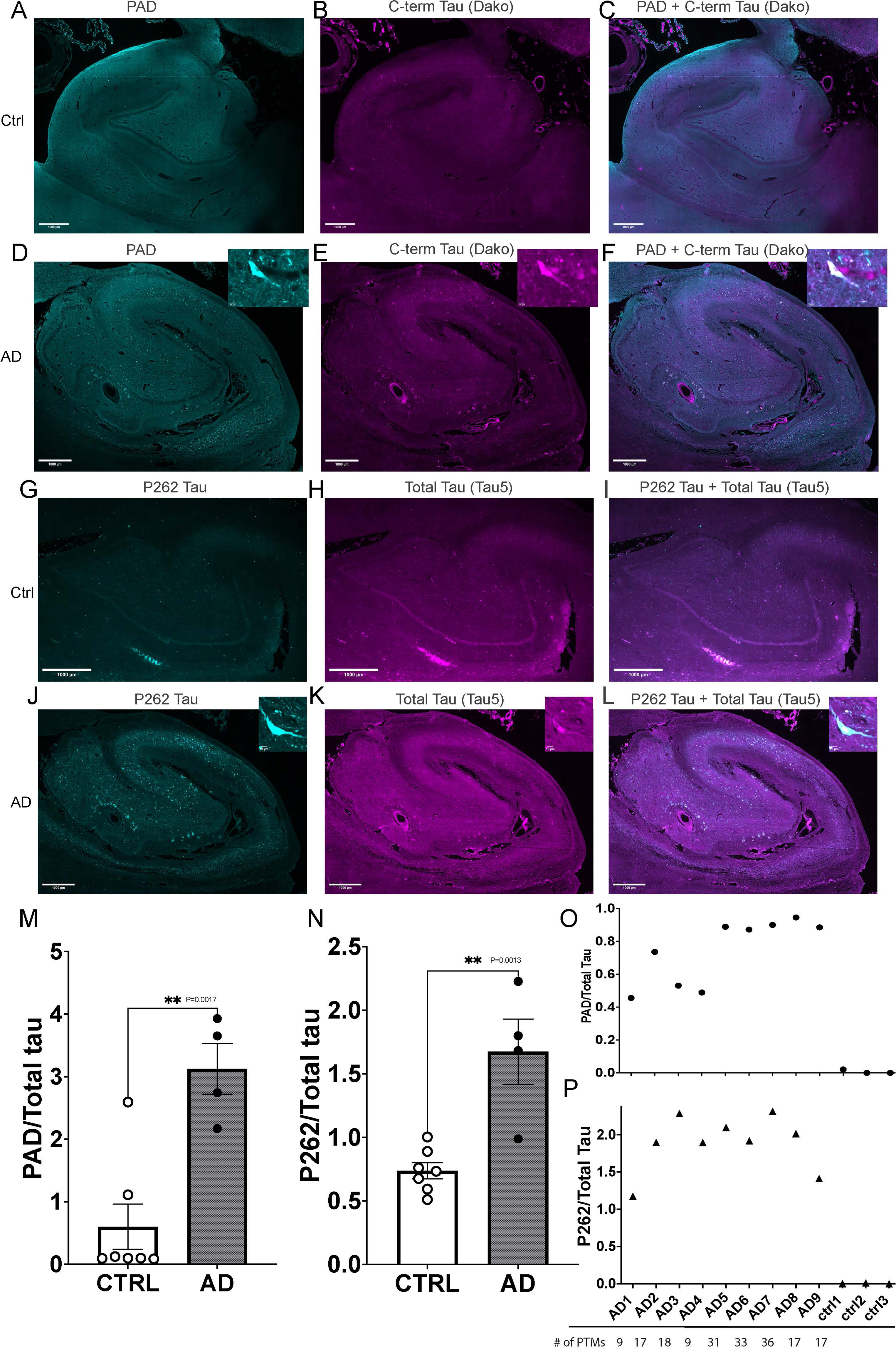
Higher PAD and p262-Tau immunoreactivities in AD patient brains. The hippocampus in age-matched controls showed low PAD (**A-C**) and p262-Tau (**G-I**) signals, while the AD hippocampus showed significantly higher PAD (**D-F**) and p262-Tau (**J-L**) signals. All data were normalized to total Tau levels and quantified (**M-N**). n=7 for controls and n=4 for AD. (**O-P**) PAD and p262-Tau immunoreactivities were much higher in high molecular weight Tau fractions from 7 AD patients, whose Tau PTMs were previously profiled by Mass Spectrometry and whose Tau seeding activities were also quantified^7^. AD 5-7, which showed the highest numbers of PTMs (including phosphorylation, ubiquitination, and acetylation) and highest seeding activities, also showed the highest PAD and p262-Tau signals. In contrast, AD1, which had the lowest PTM number and seeding activity, showed the lowest PAD and p262-Tau signals.

Altogether, these data suggest that Tau misfolding partially characterized as PAD exposure due to the loss of compact paperclip self-fold correlates with Tau pathologies in AD patients. High PAD and p262 immunoreactivities correlate with high numbers of Tau PTMs and seeding competence, suggesting the connection between Tau conformation and Tau biochemical modifications/properties contributing to Tauopathies. Therefore, these two parameters may serve as biomarkers for disease progression and responses to treatment, together with other existing Tau biomarkers. MT association is compromised in AD patients as expected, but preserved especially at earlier Braak stages, presenting another level of disease heterogeneity and challenging conventional wisdom viewing MT loss as a primary cause of neurodegeneration associated with Tauopathies. The contribution of Tau-MT interaction to disease pathology will be discussed below.

## DISCUSSION

In healthy neurons, Tau is present as soluble monomers associated with MTs. Disruptions in Tau-MT interactions promote Tau misfolding and aggregation. In AD and ADRDs, Tau undergoes aberrant auto-assembly into ordered filaments, enriched in beta-sheets. Studies of human patient brains affected by these pathologies revealed diverse filament types, each associated with specific Tau isoforms and mutations. While all filaments were enriched in PTMs, especially phosphorylations, their 3-D structures were specific to the type of Tauopathy. In addition to fibrils, various forms of misfolded Tau intermediates including monomers and oligomers can be pathogenic. However, questions remain as to which species of Tau are toxic and why. Equally puzzling are the relationships and potential transitions between various species in each disease. To effectively treat these diseases, a molecular-level understanding of the physiological Tau-MT interaction and its pathological disruption, particularly before the formation of likely irreversible tangles, seems crucial.

Here, we employed HDX-MS and mapped, at the peptide level, interactions between MTs and WT Tau or Tau variants mimicking AD-associated PTMs or ADRD-related mutation. These interactions and Tau conformations were further characterized in a cellular context using isolated axoplasm and in a disease context using THY22 mice and human AD patients. Taken together, our data show that WT Tau interacts with MTs *via* domains beyond canonical MTBRs in a continuous and cooperative fashion- “dynamic oscillation”- and that MT-bound WT Tau likely maintains a paperclip fold where N-term PAD motif is normally sequestered. In contrast, AD/ADRD-associated Tau variants have altered binding to MTs beyond the canonical MTBRs, an interaction more complex than the simple loss of binding that has long been assumed. A shared structural signature among most of these soluble monomeric Tau species associated with pathology is the exposure of PAD, which activates downstream PPI-GSK3β to inhibit axonal transport. However, conformational changes are not limited to PAD exposure as other biologically active motifs may also play a role in Tau pathology. In addition to PAD exposure, Tau phosphorylation on S262 seems to correlate with disease progression in both THY-Tau22 mouse and AD patient brains. Reduced MT binding is also observed to correlate with disease progression, especially in axonal tracts, though MT loss is unlikely to be the primary cause of the disease pathology. Finally, there is a significant heterogeneity in neuronal vulnerability regarding Tau misfolding, PTMs, and MT association, suggesting critical disease-related molecular mechanisms that may indicate early biomarkers and therapeutic targets.

### Unique Tau-MT Interactions through Sequences with Different Binding-Release Properties

Our HDX-MS experiments demonstrate that Tau alone, whether WT or mutant, behaves as an intrinsically disordered protein as evidenced by the rapid and complete deuterium exchange of its peptides, consistent with findings from other biophysical techniques, including NMR, FRET, and EPR spectroscopy^64^. However, binding of Tau to MT imposes order on specific regions of Tau that include both the canonical MTBD as well as adjacent regions and more distant domains (N-terminal and C-terminal).

To obtain meaningful HDX-MS data regarding Tau-MT interactions, we overcame a significant HDX-MS technical challenge: ensuring highly accurate and reproducible peptide data from Tau monomers in the presence of ∼10-fold molar excess of tubulin peptides; an inevitable excess due to the higher molecular weight of MT polymer bound to each Tau molecule at the physiological ratio. Upon MT binding, WT Tau could be divided into three major regions based on their protection patterns (Table 1). The region of direct interaction with MTs, we termed the Core Binding Region, was larger than the MTBRs initially described in the literature based on models derived from truncated versions of Tau^12^. HDX-MS methodology allowed us to study dynamically the full sequence of Tau and revealed that the Core Binding Region was flanked by the N- and C-terminal regions, which displayed similar protection patterns in terms of amplitude and kinetics. These three regions were connected by two hinge-like segments, which exhibited intermediate changes upon MT binding and may mediate the self-folding of Tau into paperclip conformation.

We modeled in Pymol several possibilities of N- and C- termini interactions based on the known cryoEM structures of MTBRs of Tau and MTs, as well as their interactions (Fig.S19). The classical paperclip conformation as originally proposed (S19A), where a major part of the N-terminal region is far from either MTBRs of Tau or MTs, is not consistent with the HDX data showing high protection in this region. A modified paperclip conformation (S19B) incorporates strong binding sites between regions outside the MTBRs of Tau and MTs as suggested by HDX data. Modeling could produce a parallel paperclip conformation (S19C) where N-term and C-term interact in a reversed direction to that of the classical paperclip. This allows close proximity of N-terminal regions of Tau to its MTBRs. Finally, MT-bound paperclip conformation (S19D) shows that Tau could fold into the classical paperclip while interacting with multiple adjacent MT polymers, consistent with the HDX data showing protection of almost all Tau regions in the presence of MTs.

With preassembled MTs, some Tau variants (S262E, S262A, and P301L) retained three major binding regions overlapping to some extent with WT Tau, but with different AA sequences and varied binding strengths (Table 1). The S262E mutation significantly weakened the interaction of R1 and part of R2 with MTs, but extended the Core Binding Region to R’, which belonged to Hinge Segment 2 in WT, with stronger MT interaction in part of R2, R3, R4, and R’ than WT, suggesting a shift in Core Binding Region towards the C-terminus. It is worth noting that S262E Tau still binds MTs, but this binding is likely to be transient and more prone to detachment. Together with the loss of Hinge Segment 2 and immediate and full deuteration at both N-and C-termini, this also supports our hypothesis that S262E acquires an open conformation with both termini exposed. It is worth noting that P258-266 in WT was fully deuterated after only 5 seconds and S262 fits into the junction between α and β tubulin subunits. Phosphorylation of S262 likely induces a steric clash with E434 of the C-terminus of tubulin to push this part of Tau off the MTs. Interestingly, P301L demonstrated stronger binding to MTs than WT, with the loss of both Hinge Segments as they were incorporated into the Core Binding Region, suggesting a broader MT-Tau interface, which could potentially pull the very N- and C-termini apart. The HDX profile of P301L-Tau also showed strong protection in both N- and C-termini, indicating sequestration either due to compact paperclip folding like the WT, or direct binding to MTs, or self-assembly similar to a recent model of “cohesive envelopes of Tau” formed cooperatively on MTs^61^. Puzzling enough, 2 peptides (AA 258-266 and AA 281-290) showed significantly less protection than all the other domains. This indicates that despite an overall increased protection of the MTBRs in P301L than its homologue in WT, some parts seemed to be “off” of the correct binding, suggesting that the “cooperativity” inside this domain of P301L Tau is partially lost. This also provides the possibility for S262, which is within AA258-266, to be less protected by MTs and thus more accessible for phosphorylation in FTD mice and patients with P301L/S mutations. These changes in the dynamic conformation of Tau with mutations or specific PTMs help to explain why some mutations and PTMs are pathological as well as why there are differences in different tauopathies.

Finally, HDX profiles of AD-mimic Tau proteins bearing multiple PTMs (acetylation for 9KQ, combinations of acetylation and phosphorylation for HMW and LMW) showed significant loss of MT binding and increased exposure of N- and C-termini, with HMW being most severe. HMW mimics AD Tau as identified by proteomics while LMW is also present in age-matched non-AD controls^8^. Unexpectedly, 9KQ exhibited a very different profile where several regions of Tau, including N- and C-termini and MTBRs, were relatively sequestered even in the absence of MTs and these sequestrations were enhanced by MT binding, suggesting a different conformation and binding mode that will be discussed below. All these data suggest that accumulation of PTMs associated with AD or aging can also impact Tau-MT interactions and Tau folding, consistent with the proposed distinct pathological roles of certain PTMs. Future studies will use single mutations and different combinations of mutations and PTMs to further elucidate the specific roles for these PTMs in Tau structure and function. The sequence of occurrences for these PTMs and their differential effects on Tau structure may render a certain fold more vulnerable to additional PTMs, prone to aggregation, and increase toxicity to neurons.

### Dynamic/Continuous/Cooperative Tau-MT Binding and its Disruption by AD/ADRD Tau Mutations

Deuteration profiles for Tau alone and Tau-MT complex exhibited significant differences, demonstrating that Tau is largely protected from deuteration in the presence of MTs, with the strongest sequestration seen for MTBRs and substantial, though weaker, sequestration in both N- and C-termini. However, this protection seemed transient as deuteration occurred within 3 seconds and all peptides in all regions of Tau became fully deuterated within a minute, much shorter than most complexes studied by HDX-MS (Fig.1C). This could result from complete attachment and detachment of the whole Tau molecule from MTs or from constant “binding-release” of various subdomains of Tau that move continuously and cooperatively on MTs as proposed in our “dynamic oscillation” model, similar to the “kiss-and-hop” model previously proposed based on single-molecule tracking of Tau in living neurons^65^. A complete detachment of Tau from MTs in such a short time window contradicts the experimental observations that Tau seems to be associated with MTs and promotes MT assembly *in vitro*. However, a complete and stable association of Tau with MTs also presents problems to MT tracks, which require a certain degree of dynamicity that will allow growth and rewiring as well as the maintenance of active transport of various cargoes. Therefore, we speculate that WT Tau, as a whole molecule, is associated with MTs, while different subdomains take turns to bind and release along the MTs in a continuous and cooperative manner, a model that best fits our HDX data. This dynamic model incorporating a multitude of small yet transient Tau-MT interactions keeps Tau in close vicinity to MTs and allows local movement on the MT lattice. It is also necessary for maintaining the structure and function of neuronal MT cytoskeleton as well as its interactions with other MAPs.

Modified Tau proteins with point mutants such as S262E seem to lose this dynamic but continuous association with MTs and may fall off MTs more easily and completely as one piece, as discussed earlier. The loss of MT binding leads to the loss of the compact paperclip conformation and may further facilitate misfolding and aggregation of Tau *via* MTBRs and other exposed motifs, aberrant protein-protein interactions, and downstream signaling pathways that may be toxic to neurons, as well as additional PTMs that may contribute to Tau toxicity, accumulation, and spreading, as observed in AD/ADRD mouse models and patient samples.

Altogether, our HDX data underscores the structural complexity of Tau and Tau-MT interactions and offers a higher-resolution map of interaction regions, revealing subtle differences in protection and kinetics as well as conformational dynamics. In particular, the slow deuteration of regions outside the canonical MTBRs, including parts of the proline-rich domain P2 (AA 226–241) and R’ domain (AA 370-393), highlights their critical roles in MT binding. These findings indicate a broader interaction interface than previously anticipated, which could help reconcile discrepancies observed across different experimental approaches. Critically, our HDX-MS data is in full agreement with the recent interaction models developed from molecular dynamic simulations applied to cryoEM^37^ and solid-state NMR^16^. The binding of Tau to MTs through repeated domains is also consistent with recent cryo-EM structures of other MT-associated proteins. These include CFAP21, Pierce1, and Pierce2 in doublet MTs^66^ and SPM1 and FAP363 on the inner wall of cortical MTs^67^, all of which present extended AA sequences anchored on MTs, suggesting a common mechanism for this type of interactions. Finally, HDX profiles of mutant Tau also show the existence of distinct structural rearrangements of Tau upon MT binding that can be strengthened or weakened by disease-associated modifications and may have implications for understanding the structure and function of normal Tau as well as pathological mutations.

### PAD Exposure as a Shared Signature of Several Tau Mutants and its Effects on Neuronal Function

Previously, we showed that N-term PAD motif of Tau (AA 2-18) mediated the inhibition of axonal transport induced by various pathological versions of Tau and its immunoreactivity increased in AD patient brains^29,32^. The PAD motif binds PP1 to activate GSK3β, whose activity is elevated in AD. GSK3β has many neuronal substrates, for example, it phosphorylates kinesin light chain, leading to premature release of cargoes anterogradely transported along the axons and reduced number of vesicles reaching the synapses^68^. Removing PAD from pathological Tau abolished its inhibitory effect on axonal transport and PAD motif alone was sufficient to activate PP1-GSK3β and inhibit anterograde transport^29,32^. To test whether PAD exposure is a common signature of AD/ADRD Tau species, we focused on the PAD protection in the HDX data and immunostaining of PAD in various model systems: squid axoplasm perfused with human Tau variants, THY22-Tau mouse model, and AD patients with well-characterized Tau PTMs. Our HDX data showed that S262E and HMW Tau proteins completely lost protection against HD exchange in PAD, while PAD in P301L, 9KQ, and LMW Tau proteins maintained some protection to various degrees. Unexpectedly, PAD seemed sequestered in 9KQ Tau and to a lesser extent in LMW, even in the absence of MTs, suggesting intra- or inter-molecular interactions between PAD and other parts of Tau. 9KQ and LMW Tau proteins share the same pseudoacetylation K353Q, while 9KQ carries additional 8 other pseudoacetylations and LMW Tau carries additional 8 pseudophosphorylations including T181D, S199D, T217D, T231D, T403D, and S404D, which are also present in HMW Tau. Interestingly, pseudophosphorylation on S262 is only on S262E and HMW, but not the others, suggesting a critical role of this phosphorylation in Tau folding and interaction with MTs, which may explain the observation that S262 phosphorylation level correlated with AD progression in clinic^11^.

The differences in the HDX profile of PAD are intriguing, so we carefully examined the same Tau proteins in the context of squid axoplasm with intact endogenous axonal MTs using TNT1 antibody specifically recognizing part of PAD using IHC. TNT1 antibody shows low immunoreactivity when PAD is sequestered in a paperclip fold of WT Tau and increases when Tau opens to expose the N-terminus. Interestingly, all Tau variants showed increased PAD immunoreactivity with S262E and HMW having the highest signal, suggesting the loss of sequestration as indicated by HDX-MS. However, P301L, 9KQ, and LMW also showed an intermediate increase in PAD immunoreactivity, suggesting a partial loss of PAD sequestration. Interestingly, except for S262E and HMW that showed significant loss of MT binding, P301L, 9KQ, and LMW were still associated with MTs, like WT. Therefore, we speculate that phosphorylation on S262 in S262E and HMW mutants reduces their interactions with MTs leading to the exposure of PAD, as MT binding keeps WT Tau in a folded paperclip conformation to sequester PAD. P301L, though tightly associated with MTs as suggested by both HDX-MS and immunostaining data, may adopt an open conformation where PAD is exposed but may be sequestered once interacting with other neighboring Tau molecules to form an envelope surrounding MTs, especially at saturating concentrations as used in HDX-MS. Surprisingly, 9KQ Tau uniquely induced local rearrangement of MT cytoskeleton to create empty circles that spaced out parts of neighboring MT arrays. In the light of the HDX data suggesting potential inter- or intra-molecular binding of 9KQ Tau even in the absence of MTs, we predict that 9KQ folds into a circular structure within itself and forces its bound MTs (mainly *via* R1, as the rest of MTBRs of 9KQ showed reduced MT binding similar to HMW Tau, Fig. 2I) into these structures. Future studies will use single pseudoacetylations and various combinations of these single mutations to explore the underlying molecular mechanisms and model this unique behavior. Finally, LMW presents an intermediate behavior regarding its MT binding and PAD exposure, as LMW shares some AD-related PTMs but is also present in age-matched AD control patients. We conclude that LMW has the tendency to fall off MTs and misfold, susceptible to further PTMs, and may share a certain level of pathogenicity as AD-HMW Tau.

To further confirm the exposure of PAD and explore the potential toxicity of these Tau variants in neuronal function, we measured vesicular axonal trafficking, a sensitive readout of axonal and synaptic function, in the axoplasm perfused with these Tau proteins. S262E, 9KQ, HMW, LMW, and P301L all inhibited anterograde transport as PAD alone did, and this inhibition was prevented by TNT1 antibody that sequesters PAD, consistent with the structural prediction of PAD exposure in these proteins. However, some Tau variants also inhibited retrograde transport to various degrees, different from the action of PAD alone. Though, inhibition of retrograde transport can result from an overall inhibition of anterograde transport, TNT1 failed to fully prevent retrograde transport defects induced by P301L and HMW, suggesting additional mechanisms that may inhibit dynein motility and/or its binding to cargoes, which are normally transported retrogradely.

Altogether, these data suggest that PAD exposure is a common feature of various AD/ADRD-Tau variants, indicating Tau misfolding and toxicity in neurons, especially in the context of axonal transport of synaptic vesicles, which are critical for normal axonal maintenance and synaptic function. However, disease associated Tau variants are likely to acquire additional misfolds that disrupt normal binding partners such as MTs, induce abnormal protein-protein interactions, and activate aberrant signaling pathways, leading to additional toxicity. HDX measurement, immunostaining pattern, and axonal transport assay also jointly indicate that phosphorylation on S262 may play a major role in Tau folding, MT binding, and axonal function. Thus, we further examined this PTM in mouse and human AD/ADRD models as discussed below.

### The Role of Tau S262 Phosphorylation (p262 Tau) in AD/ADRD Disease Progression

There are numerous hyperphosphorylation sites on Tau in AD, some of which (e.g., pTau181, pTau217, and pTau231) serve as biomarkers as they are associated with *in vivo* and autopsy-verified diagnosis in AD^69^. As Tau is an abundant protein in neurons and has more than 80 known phosphorylation sites, it is likely to serve as an ideal substrate for multiple kinases aberrantly activated in AD. It is still unclear as to how or whether different Tau phosphorylations drive disease progression.

Our HDX data suggested that S262 sits in the conjunction between α and β tubulin subunits and its phosphorylation induces a steric clash to interfere with MT binding. Our axonal transport data suggested that pseudophosphorylated Tau on S262 and Tau phosphorylated on S262 by MARK kinase *in vitro* both inhibit axonal transport, which was dependent on exposure of the N-term PAD in p262 Tau. In the THY22 mouse model bearing FTD Tau mutations, p262 Tau immunoreactivity is much higher when compared to WT and increases as the pathology progresses during aging. PAD immunoreactivity is significantly higher than in WT but seems to saturate at an early stage (i.e., 3mo) without further increases in older mice, indicating chronic exposure in these FTD-mutant Tau proteins. This suggests that mutant Tau with an open structure is more likely to be phosphorylated by multiple kinases including MARK, whose activity increases as pathology progresses.

Interestingly, p262 Tau correlates with AD progression in AD patients and seems more abundant in AD Tau fractions that are enriched in seeding competent Tau with the most PTMs especially phosphorylation. P262 Tau is more likely to detach from MTs due to the loss of cooperative binding and thus is prone to further misfolding and aggregating. Given the toxic effects of p262 Tau on axonal transport, this is likely to be a direct cause of axonal and synaptic dysfunction in affected neurons in AD/ADRDs.

However, loss of MT binding is not required for toxicity. P301L/S Tau remains associated with MTs based on both HDX data, immunostaining data in isolated axoplasm perfused with P301L Tau and in THY22 mice at early stages (3-6mo), when PAD immunoreactivity is elevated. This supports the previous speculation that these mutant Tau proteins have an intrinsically open structure even when bound to MTs. Considering the HDX data demonstrating tight association between P301L-Tau and MTs, this also challenges a major hypothesis in the field that disease associated Tau causes neurodegeneration due to its loss of function as it no longer binds and stabilizes MTs.

Altogether, disease associated mutant Tau proteins seem more prone to hyperphosphorylation, perhaps due to misfolded and open Tau being both more susceptible to kinases and more likely to activate downstream signaling pathways. Among these phosphorylations, p262 is correlated with disease progression, toxic to axonal transport, and likely to detach from MTs abruptly to allow further misfolding and aggregation. This makes p262 Tau a potential biomarker and driver of neuronal dysfunction and Tau pathology during AD progression.

### The complex role of Tau-MT binding in AD/ADRDs

Both loss of function of Tau leading to MT destabilization and toxic gain of function for pathological Tau have been proposed and debated as a primary disease mechanism. We previously established that axonal MTs are uniquely stable due to many factors other than Tau, such as unique PTMs and various MAPs^26,27^. Recent studies also suggest that Tau regulates MT dynamics *via* labile domains^13,70,71^ rather than stabilizing MTs. Finally, Tau KO mice don’t recapitulate AD phenotypes and MT stability in those mice are largely normal^72^. Our data demonstrated that P301L Tau was associated with MTs *in vitro* and *in vivo* and the MT structures were intact initially. In contrast, our previous and current data showed that various pathological Tau proteins at 2µM inhibit axonal transport without affecting MT structures, suggesting a toxic gain of function. Therefore, it is unlikely that loss of MT binding and failure in stabilizing MTs by disease-associated Tau is a primary cause of neurodegeneration in AD/ADRDs. Does Tau-MT binding play a role in Tauopathies?

Our HDX data indicates that MT binding keeps Tau folded into a compact paperclip conformation. AD/ADRD-related phosphorylations and mutations either reduce MT binding leading to an open conformation of Tau (e.g., in the case of p262 Tau) or directly induce an open conformation of Tau bound to MTs leading to more PTMs and eventually detach Tau from MTs for further aggregation (e.g., in the case of P301L Tau). This does not exclude the possibility that impairment of axonal transport and additional PTMs as a result of toxic gain of disease-modified Tau function may eventually lead to altered MT structures and functions, as evidenced by our SIM data showing reductions in Tau-MT association in axons from THY22 mice (after 6mo) and in postmortem AD patient tissues. Furthermore, some PTMs of Tau could also change MT spacing and rearrange MT arrays as suggested by local effects on MT structure with 9KQ Tau. Finally, some disease-associated PTMs, such as acetylation, may differentially affect both MTs and Tau. Experiments manipulating enzymes responsible for these PTMs by either genetic or pharmacological methods should be analyzed carefully in the context of all cytoskeletal substrates and the specificity of the action on the proposed target.

An unexpected finding from our HDX data is that Taxol-induced MT conformational changes caused a global weakening of Tau-MT affinity while exhibiting regional specificity in its impact. Taxol was initially included to exclude potential effects of MT depolymerization during HDX experiments that could alter rates of deuteration over time. HDX data showed that Taxol did not reduce Tau deuteration, but Taxol-stabilized MTs showed lower affinity for Tau in all conventional MTBRs. This effect was partially offset by an increased binding to R’ domain, highlighting the dynamic nature of MT-Tau interaction. This suggests that Taxol could potentially exacerbate Tau misfolding and aggregation by reducing normal interactions with MTs. Further investigation is needed to fully understand the implications of Taxol and similar MT stabilizing drugs for Tau function and their proposed use in neurological conditions such as Tauopathies and axonal regeneration.

Altogether, our data suggest that while loss of MT binding of Tau is unlikely to induce MT destabilization but may contribute to Tau misfolding and aggregation *via* exposed MTBRs that are no longer sequestered by MTs and/or N/C-termini that lose their interactions in the paperclip conformation. This secondary effect may exacerbate disease pathogenesis especially at the late stage. Further, misfolded Tau may affect MT-mediated axonal trafficking, which will in turn affect MT structure and function, as observed in both animal models and postmortem patient tissues.

### Tau as an Intrinsically Disordered Protein

The question of how to interpret the HDX-MS results in the context of Tau as an IDP was an initial concern. We took advantage of existing data available for WT-Tau fragments bound to MTs from high resolution methods (mainly cryoEM and NMR) and assigned the relative uptake for the domains for which such data is available (e.g., MTBRs). However, structural data are limited for full length Tau. All Tau proteins used in this study are based on 2N4R isoform and the effects of mutations or PTMs on different Tau isoforms may differ^73^. Furthermore. the MTs used *in vitro* studies were purified from young pig brains, while MTs from different ages may have different PTMs during aging and tubulin isoform composition.

Our data from the isolated axoplasm, mouse models, and human tissues, partially mitigate this concern, but future studies may need to address this heterogeneity in both Tau and MTs. This would include MTs purified from aged mouse and human AD brains to further examine the interactions between Tau and MTs in the context of aging and neurodegeneration. HDX-MS also has the potential to probe endogenous Tau proteins in longitudinal samples from AD models such as mouse brains at various ages and patient biofluids at various disease stages. Though our data suggest that Tau binds MTs in a continuous, dynamic and cooperative manner, and may not completely fall off MTs, we should keep in mind that neuronal MTs are often compact in healthy axons, thus even if Tau is fully detached from one MT polymer, it may soon bind a neighboring MT polymer like molecular motors. However, this rebinding opportunity may be significantly compromised by reduced MT density in aged and degenerating neurons.

Finally, our data suggest conformational changes of soluble monomeric Tau due to mutations and PTMs. These misfolded monomers may assemble into higher order oligomers and insoluble fibrils *via* exposed MTBRs that are no longer sequestered by MT binding or exposed N and C termini that are no longer sequestered by the paperclip fold. This progression is likely to be heterogeneous, as fibrils in different Tauopathies acquire unique folds and this transition needs to be studied more thoroughly and dynamically in the future. Understanding both the shared and unique conformational signatures and whether and how they transition in between will help us understand the role of Tau in AD/ADRD pathogenesis.

### Statistical Information

All experiments were repeated at least five times. The data were analyzed by one-way ANOVA followed by the Tukey *post-hoc* test (or nonparametric multiple t-tests, without assuming consistent SD) and plotted in Prism 7 (GraphPad software). Quantitative data were plotted as mean ± SEM. P values were calculated, and four statistical thresholds were marked: P<0.00001, P<0.0001, P<0.001, and P<0.005 (P<0.05 indicated statistical significance).

## Data availability

The authors confirm that the data supporting the findings of this study are available within the article and its supplementary material.

## Author Contribution

YS and FG conceived the ideas, designed the study, and wrote the manuscript. FG and YS performed all HDX-MS experiments and data analyses. SO, TB, and YS performed all mouse and human studies and related image analyses. JRD, CD, ZF, and YS designed constructs, purified, tagged, and characterized all the Tau proteins used in this study. NQ and YS prepared all human brain lysates, performed dotblots, and analyzed data. SO, STB, and YS performed all squid studies and analyzed data. BH and CB provided invaluable suggestions and resources throughout the study. All authors provided input to the manuscript.

## Acknowledgments

We dedicate this manuscript to Lester (Skip) Binder and David Remsen, who passed away but continue to live through our Tau studies and squid experiments, respectively. We thank John Engen for suggestions and resources regarding HDX-MS experiments, Nick Kannan for the TNT1 antibody and helpful discussions about Tau in AD in the last 10+ years, and Gerardo Morfini for his help with squid axoplasm studies at MBL and his scientific friendship over the years. We acknowledge fruitful discussions with Alain Dautant regarding Tau structures and with Tyler Levy regarding data presentation. We also thank Drs. Merit Cudkowicz, Craig Blackstone, and Bradley Hyman for fully supporting the Song Laboratory at MGH.

## Funding

The authors would like to acknowledge all the funding resources: NIH grants [P30AG062421 (Developmental Project) and AG072516], a Jack Satter Foundation grant, a Pape Adams ALS Transformative Scholar Award, an AARG grant from the Alzheimer’s Association, and Healey Center ALS Awards from MGH to YS. Additional support was provided by NIH grants R01NS023868, R21 AG067772 and a Zenith Award from the Alzheimer’s Association to STB, as well as P30AG062421 for Massachusetts Alzheimer Disease Research Center and Rainwater Foundation to BTH, and Jack Satter Award to JRD.

## Competing interests

The authors report no competing interests.

*This manuscript is under consideration at a journal.

## Notes

### Competing Interest Statement

The authors have declared no competing interest.

